# Cell-specific and targeted delivery of RNA moieties

**DOI:** 10.1101/2020.01.03.893818

**Authors:** Aditi Bhargava, Peter Ohara, Luc Jasmin

## Abstract

Delivery of therapeutic moieties to specific cell types, such as neurons remains a challenge. Genes present in neurons are also expressed in non-neuronal cell types such as glia where they mediate non-targeted related functions. Thus, non-specific targeting of these proteins/channels has numerous unwanted side effects, as is the case with current small molecules or drug therapies. Current methodologies that use nanoparticles, lipid-mediated uptake, or mannitol in conjunction with lipids to deliver double-stranded RNA (dsRNA) have yielded mixed and unreliable results. We used a neuroanatomical tracer (B subunit of Cholera Toxin (CTB)) that binds to the ganglioside receptors (GM1) expressed on cells, including primary sensory neurons to deliver encapsulated dsRNA. This approach greatly improved delivery of dsRNA to the desired cells by enhancing uptake, reducing vehicle-mediated toxicity and protecting nucleotides from degradation by endonucleases. The delivery complex is internalized, and once inside the cell, the dsRNA naturally dissociates itself from the carrier complex and is very effective in knocking down cognate targets, both *in vivo* and *in vitro*. Past methods have used CTB-fusion proteins or chemically modified oligos or DNA moieties that have been covalently conjugated to CTB. Furthermore, CTB conjugated to an antigen, protein, or chemically modified nucleic acid is a potent activator of immune cell (T and B cells, macrophages) response, whereas CTB admixed with antigens or unmodified nucleic acids does not evoke this immune response. Importantly, in our method, the nucleic acids are *not covalently linked* to the carrier molecules. Thus, our method holds strong potential for targeted delivery of therapeutic moieties for cell types expressing GM1 receptors, including neuronal cell types.

## Introduction

Targeted delivery of a therapeutic molecule to specific cells is highly desirable due to the associated advantages. For example, the targeted delivery of the therapeutic molecule might reduce systemic side effects resulting from off target effect of the therapeutic molecule as well as immune response to the therapeutic molecule. Additionally, if a therapeutic molecule can be targeted to a particular cell where the activity of the therapeutic molecule is needed, a lower amount of the therapeutic molecule could be used, thereby lowering the cost of the treatment.

Although other methods for targeted delivery of therapeutic molecules have been proposed [1], there is still a need for specific complexes that provide effective delivery of the therapeutic molecule. The lack of methods for targeted delivery of nucleic acids *in vitro* and *in vivo* is a limiting factor in the development of localized treatment. Gene therapy mediated by RNA interference (RNAi) faces <u>three</u> major challenges: (1) delivery, (2) off-target and/or immune effects, and (3) stability and efficacy of small interfering RNAs (siRNAs). To address these challenges, we have developed a method for targeted delivery of nucleic acids (DNA, RNA) to mammalian cells, specifically neurons. This method involves encapsulating unmodified nucleic acids and linking the resultant nanoparticles to carrier molecules (e.g., proteins or glycoproteins). This approach is different from current delivery methods such as lipofectamine or other lipid-based systems [2, 3], which lack cell selectivity because they distribute the nucleic acids to all cell types. Our method delivers nucleic acids to a subset of cells in a selective and reproducible manner. This cell selectivity is possible because the carrier is cell-specific. The targeted cells internalize the carrier-nucleic acid complex and once inside the cells, the nucleic acids dissociate from the carrier complex and perform their expected biologic activity.

Currently, two forms of RNA are widely considered as candidates for RNAi. One is micro RNAs (miRs), a naturally occurring class of small non-coding RNAs that have imperfect homology to target mRNAs and usually regulate expression of many targets, rendering their use for specific gene silencing challenging. The other form is small interfering RNAs (siRNAs), which are synthetic double-stranded RNAs (dsRNAs) introduced to bypass the first few steps of RNAi (supposedly to avoid immune activation), and are incorporated into the RNA-induced silencing complex (RISC) to bring about gene silencing[4]. While siRNAs can be effective and potent in silencing gene function, they can also evoke immune responses[5]. Several siRNA sequences for a target mRNA need to be tested to confirm effective protein knockdown, and siRNAs can often degrade cognate mRNA without affecting protein expression [6]. siRNAs have been chemically modified (with 2’ fluoro or 2’ O methyl) to increase stability [7], but these siRNAs are often less effective or have side effects [8]. To further complicate their functional significance, siRNAs were recently shown to possess activation function (RNAa) in addition to their well-known suppressor function [9].

A third form of RNAi, long dsRNA (LdsRNA) has been largely overlooked. Complementary, long, antisense RNAs transcribed from the non-coding strand occur naturally in many mammalian cell types, yet their function is poorly understood. We postulate that these naturally occurring antisense transcripts can pair with their sense mRNA, forming LdsRNAs that serve as endogenous substrates for *in vivo* RNAi to regulate gene expression and function. Others and we have shown that LdsRNA works with equal, if not better potency than siRNA *in vivo* in mammalian cells [10–16]. This finding questions the dogma that LdsRNA only works in invertebrates such as worms and flies [17], and because it uses an endogenous mechanism, has the advantage of fewer off-target effects and less immune activation. Moreover, because the LdsRNAs that we use are 300-500bp long (displaying 100% sequence identity with the target mRNA), they potentially yield numerous siRNAs after dicer cleavage. Thus, designing multiple LdsRNAs is not required, and LdsRNAs have the potential to overcome most of the shortcomings of siRNAs and advance the field of RNAi-mediated therapy.

The therapeutic areas that can be targeted with this delivery method, could include any nervous system related disease. Additionally, since cholera toxin is a gut pathogen and enters the gut via epithelial cells expressing the cognate GM1 receptor via its non-toxin subunit B (CTB), RNAi encapsulated in CTB can be targeted for gastrointestinal disorders.

## Materials and methods

### Animals

Adult male Sprague-Dawley rats weighing 250-280 grams were used for isolation of dorsal root ganglia (DRG) after intrathecal injection. All animals were housed on a 12-hour light–dark cycle and had *ad libitum* access to food and water. The UCSF Institutional Animal Care and Use Committee approved all protocols used in this study.

### Materials

Cholera toxin B subunit was purchased from Sigma (Cat # C9903, St. Louis, MO). PEG-maleimide was purchased from JenKem Technology, USA (Plano, TX).

### dsRNA synthesis

cDNA of genes of interest were generated by reverse transcription of 1 μg of total RNA followed by a 30-cycle PCR using gene-specific primers. These cDNAs were then cloned into a pTOPO vector (Invitrogen, Carlsbad, CA) and sequenced to confirm identity. The forward and reverse primer sequences used to make P2X3R and NR2B dsRNA were as follows: P2X3R Forward primer: 5’ CACCTACGAGACTACCAAGTC 3’ and Reverse primer 5’ CTCAGCCTCCATCATGATAGG 3’ corresponding to nucleotides 205-688 (488 bp, GenBank accession number NM_031075), annealing temperature of 61°C. NR2B (GenBank accession number XM_017592439) Forward primer: 5’ GCTACAACACCCACGAGAAGAG 3’ and Reverse primer: 5’ GAGAGGGTCCACGCTTTCC 3’ corresponding to nucleotides 1760-2073 (313 bp) and annealing temperature of 65°C. Sense and antisense RNA were synthesized from cDNA inserts by using MegaScript RNA kit (Ambion, Austin, TX) according to the specification of the manufacturer and as previously described [11].

### Encapsulation of dsRNA and formation of complex Q (CQ)

The complex was generated in two separate steps. In the first step, the dsRNA (2-30μg) was mixed with the PEG-linker moiety (100-150mg) in a solution containing 0.2M NaCl, pH 6.8 at room temperature for 2 hours. This step allows dsRNA to be encapsulated within PEG linker. Since dsRNA does not have any phosphorothioate (Sulphur) modification, it cannot be covalently linked with meleimide groups on PEG. In the second step, the linker-dsRNA is incubated with CTB in 0.2M NaCl, pH7.0. This allows for CTB to become conjugated to the maleimide on PEG linker, resulting in CQ-LdsRNA (Fig. 1). A reaction between PEG-maleimide and CTB (CQ shell) is referred to as the carrier.

**Figure 1.**
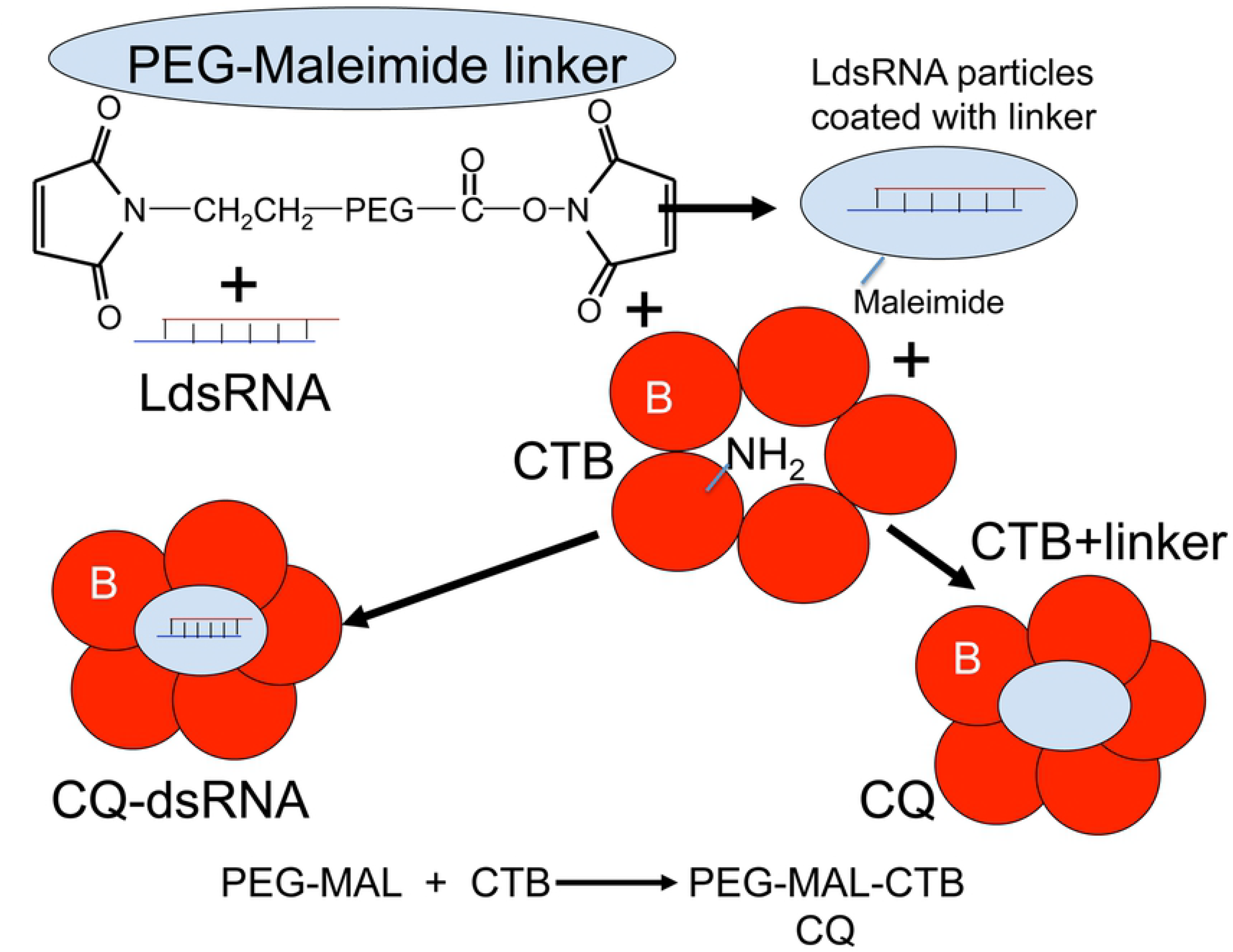
legend.Schematic showing concept for encapsulation of dsRNA inside CTB without covalent linkage. In step 1, LdsRNA is incubated with PEG-meleimide in specific salt concentration and pH, resulting in coating of dsRNA moieties with PEG-meleimide. In step 2, PEG-Maleimide coated dsRNA is incubated with CTB to give CQ-dsRNA complex. CQ alone is obtained by incubating CTB with PEG-Maleimide in presence of 0.2M NaCl, pH 6.8.

### Cell lines and cell culture

Neuro2A mouse neuroblastoma cells were obtained from ATCC (CCL131). Neuro2A cells were grown in Eagle’s Minimum Essential Medium (MEM) supplemented with 10% fetal bovine serum (FBS) and incubated at 37°C in 5% CO_2_. Cells were seeded on coverslips and differentiated by reducing serum concentration to 1% FBS for 16-20 hours followed by serum starvation for 6 hours. Subsequently, cells were maintained in MEM with 5% FBS. Differentiated Neuro2A cells were incubated with CQ-dsRNA (10μg) in 2 mL of culture medium with 5% FBS for two more days. Cells were washed, fixed with 4% paraformaldehyde and processed for immunofluorescence.

### Intrathecal injection of CQ-dsP2X3R

DsP2X3R (10μg) was encapsulated within CQ and 10 μL of CQ-dsP2X3R was injected intrathecally in the lumbar vertebrae (L4). Five days later, rats were deeply anesthetized transcardially perfused with 4% paraformaldehyde. Dorsal root ganglia corresponding to lumbar region L1-L4 were isolated and used for sectioning and immunostaining.

### Immunofluorescence and microscopy

Fifty micrometer-thick DRG sections were cut on a freezing microtome and immunofluorescence was performed on DRG sections and Neuro2A cell as described previously [18] using antibody dilutions optimized in the lab. Primary antisera directed against the following: CTB (Goat anti-CTB, dilution 1:3000, List Biologic, Campbell, CA), P2X3R (Guinea pig anti-P2X3R, dilution 1:4000, Neuromics, Edina, MN), β III tubulin (anti-mouse, dilution 1:20,000, Promega, Madison, WI), and NR2B (anti-rabbit, dilution 1:1000, Millipore Sigma).

## Results

### Formation of CQ-dsRNA complex

Modification of either CTB or dsRNAs (siRNA, LdsRNA or miRNA) modifies their properties compromises function and is immunogenic. To avoid modification steps, we first incubated LdsRNA with PEG-maleimide linker that resulted in formation of LdsRNA coated with linker, but not covalently linked [19] as the RNA could be separated from the linker on a native polyacrylamide gel by electrophoresis (Fig. 2A). In the second step of this reaction, we utilized the maleimide group on PEG to link it to the cysteine residues that served as the NH_2_ donor in CTB. The reaction is pH-dependent and takes about 40 min of incubation at 25°C. Once CTB reacts with maleimide groups, a shift in electromobility on a native PAGE can be seen between CQ-LdsRNA vs. CQ alone, with the former migrating slower than the latter (Fig. 2B). dsRNA-PEG linker complexes can be made up to one month in advance and the CQ-LdsRNA complex is stable for at least a week at 4°C.

**Figure 2.**
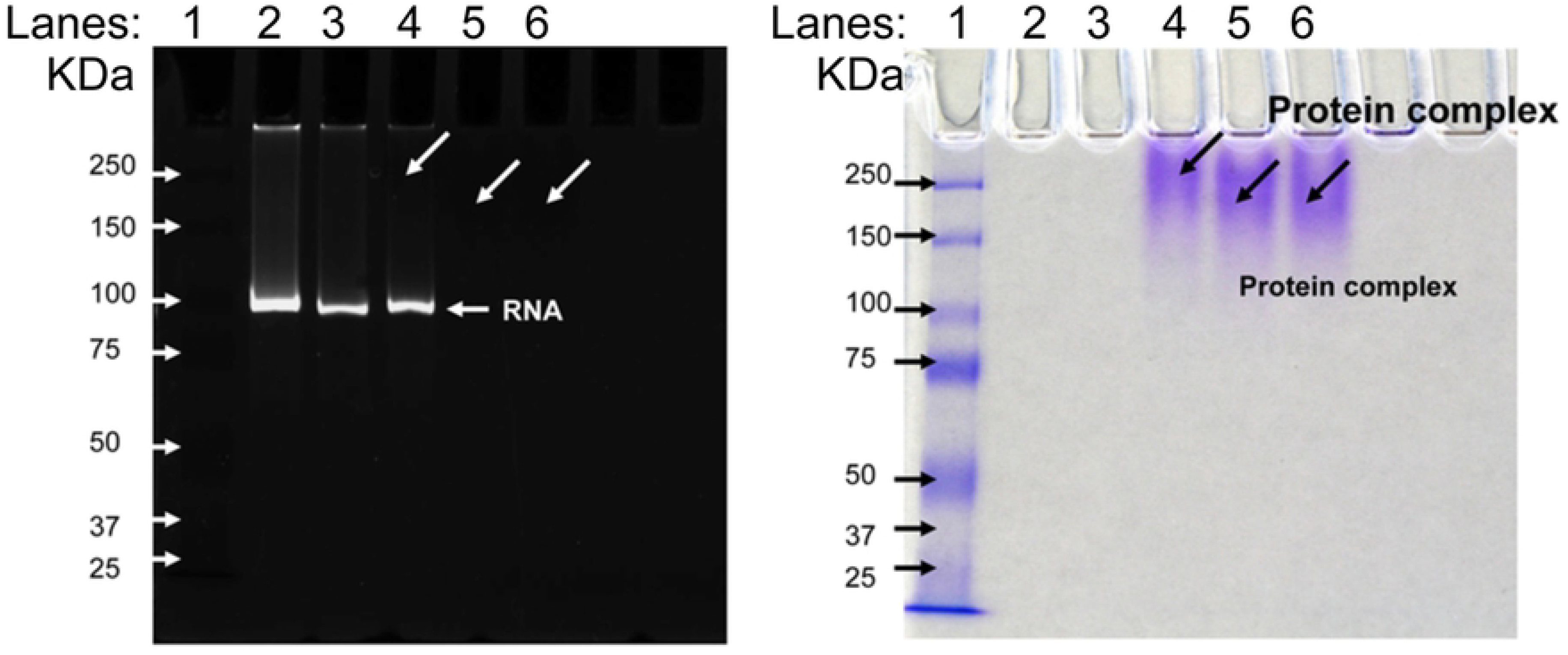
legend. dsRNA in CQ-complex is not covalently linked to PEG-meleimide. CQ-dsRNA for P2X3 receptor along with dsP2X3R-PEG-Meleimide, CTB-PEG meleimide, or CTB were electrophoresed on a 7% Native polyacrylamide gel. (A) Gel was stained with ethidium bromide to visualize dsRNA. (B) Gel was stained with Coomassie blue to visualize CQ protein complex. Lanes: 1: Marker, 2: dsP2X3R, 3: dsP2X3R-PEG-meleimide, 4: CQ-dsP2X3R, 5: CTB alone, 6: CQ (CTB-PEG-meleimide).

### CQ-dsRNA can be used *in vivo* to target specific neuronal populations in dorsal root ganglion

To test whether the CQ-dsRNA complex is viable as an *in-vivo* delivery method, we used CQ-dsP2X3R dsRNA to test efficacy of delivery and knockdown of P2X3R expression in dorsal root ganglion (DRGs). P2X3R is a purinergic receptor found on primary sensory neurons of DRGs. We injected 10μl of a CQ-dsP2X3R complex intrathecally around the lumbar spinal cord of adult rats (Fig. 3). Five days later, the rats were euthanized and the DRGs from the lumbar spinal nerves were examined using immunocytochemistry. As seen in Fig. 3 from a lumbar dorsal root ganglion, many DRG neurons contained CTB and many DRG neurons expressed P2X3R immunoreactivity (Fig. 3). However, cells containing CTB expressed no, or very reduced P2X3R immunoreactivity (Fig. 3, arrows), indicating that CQ-mediated delivery of dsP2X3R to the neurons is a viable option and efficacious in knocking down expression of the P2X3R protein in subpopulations of neurons.

**Figure 3.**
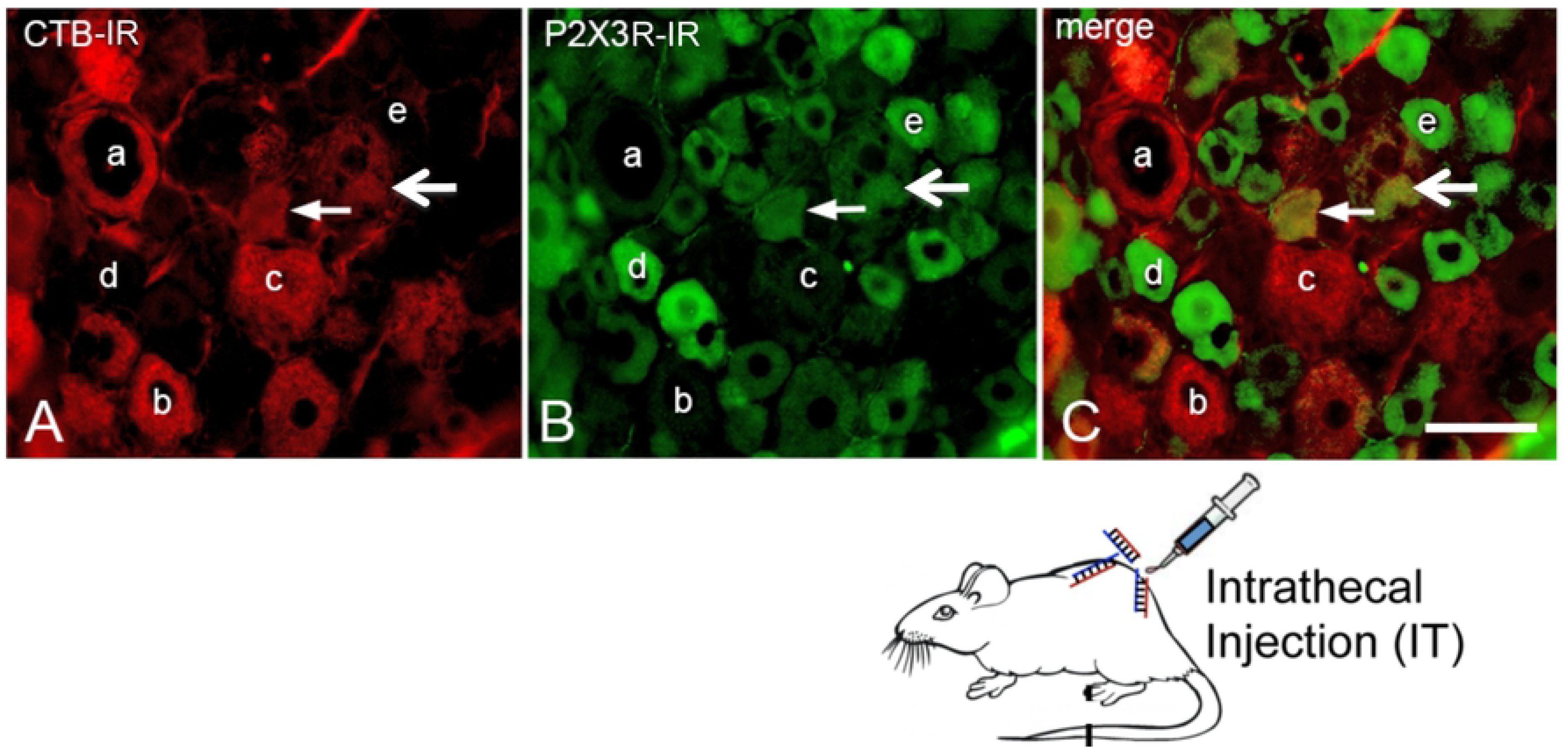
legend.CQ-dsP2X3R uptake by dorsal root ganglion neurons. Intrathecal injection of CQ-dsP2X3R resulted in uptake of the complex in large and medium diameter neurons as reflected by presence of CTB immunoreactivity (CTB-IR) in subsets of DRG neurons. a, b, c, are examples of neurons that contain CTB but show no P2X3R expression. Some neurons (e.g. arrow) are CTB positive and show reduced expression of P2X3R immunoreactivity. Neurons that strongly express P2X3R (e.g. d, e) do not contain CTB. Scale bar = 30 mm.

### Carrier or CQ by itself does not alter protein expression *in vitro*

Next, we tested whether neurons in culture can take up CQ and that CQ (carrier) itself does not alter expression of target receptors by examining the NMDA receptor subunit NR2B expression in Neuro2A cells. Immunofluorescence shows that Neuro2A cells differentiated into neuronal phenotype to project neurites and express β III tubulin immunoreactivity and co-express NR2B (Fig. 4A). CQ carrier alone added to the culture medium is taken up by efficiently by differentiated Neuro2A cells (Fig. 4B, CTB immunoreactivity) and does not alter the expression of NR2B.

**Figure 4.**
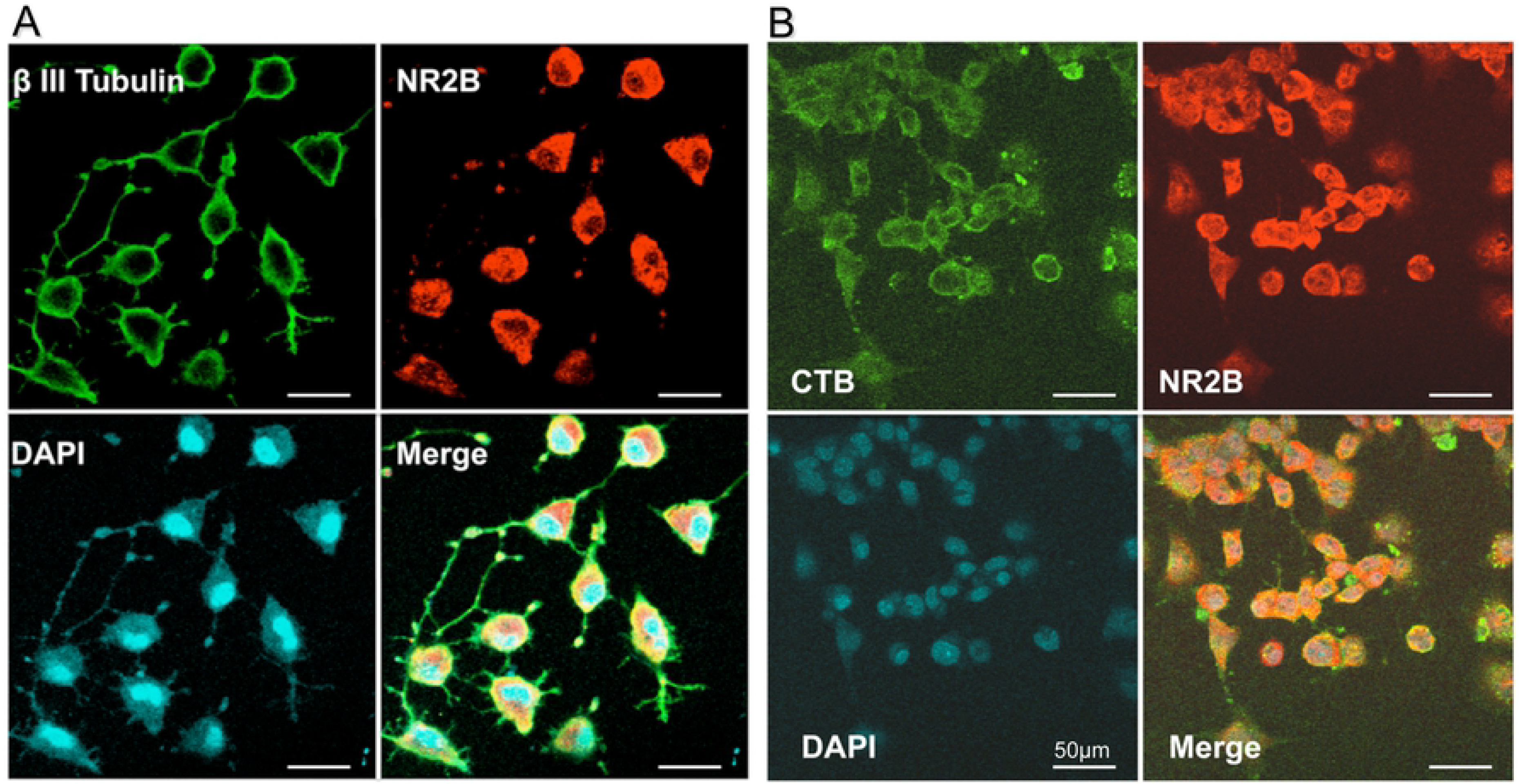
legend. Differentiated Neuro2A cells take up CQ-dsRNA complex. (A) Neuro2A cells can be differentiated and expression β III tubulin, a marker for neurons. In addition, all Neuro2A cells express NR2B subunit and co-localize with β III tubulin (merge). (B) Incubation of CQ in culture medium resulted in uptake of CTB by Neuro2A cells, but did not affect NR2B expression, suggesting that CTB-PEG meleimide by itself does not alter expression of proteins. Scale bar: 50μm.

### *In vitro* uptake of CQ-dsRNA by neuronal cell line Neuro2A

Since all Neuro2A cells express NR2B, if CQ-dsNR2B is selectively taken up by a subset of neurons, we should expect knockdown of NR2B in those subsets of neurons that take up CQ complex. Cells were incubated with CQ-dsNR2B complex for 7-24 hours, and then transferred to normal culture medium without CQ-dsNR2B. Seven hours after incubation with CQ-dsNR2B, uptake was evident, but no knockdown of NR2B immunoreactivity was evident at this short time point (Fig. 5A). Two days after incubation with CQ-dsNR2B complex, Neuro2A cells showed knockdown of NR2B in all cells that were positive for CTB immunoreactivity (Fig. 5B). In cells that did not take up CQ-dsNR2B, immunoreactive NR2B was clearly visible (Fig. 5B merge).

**Figure 5.**
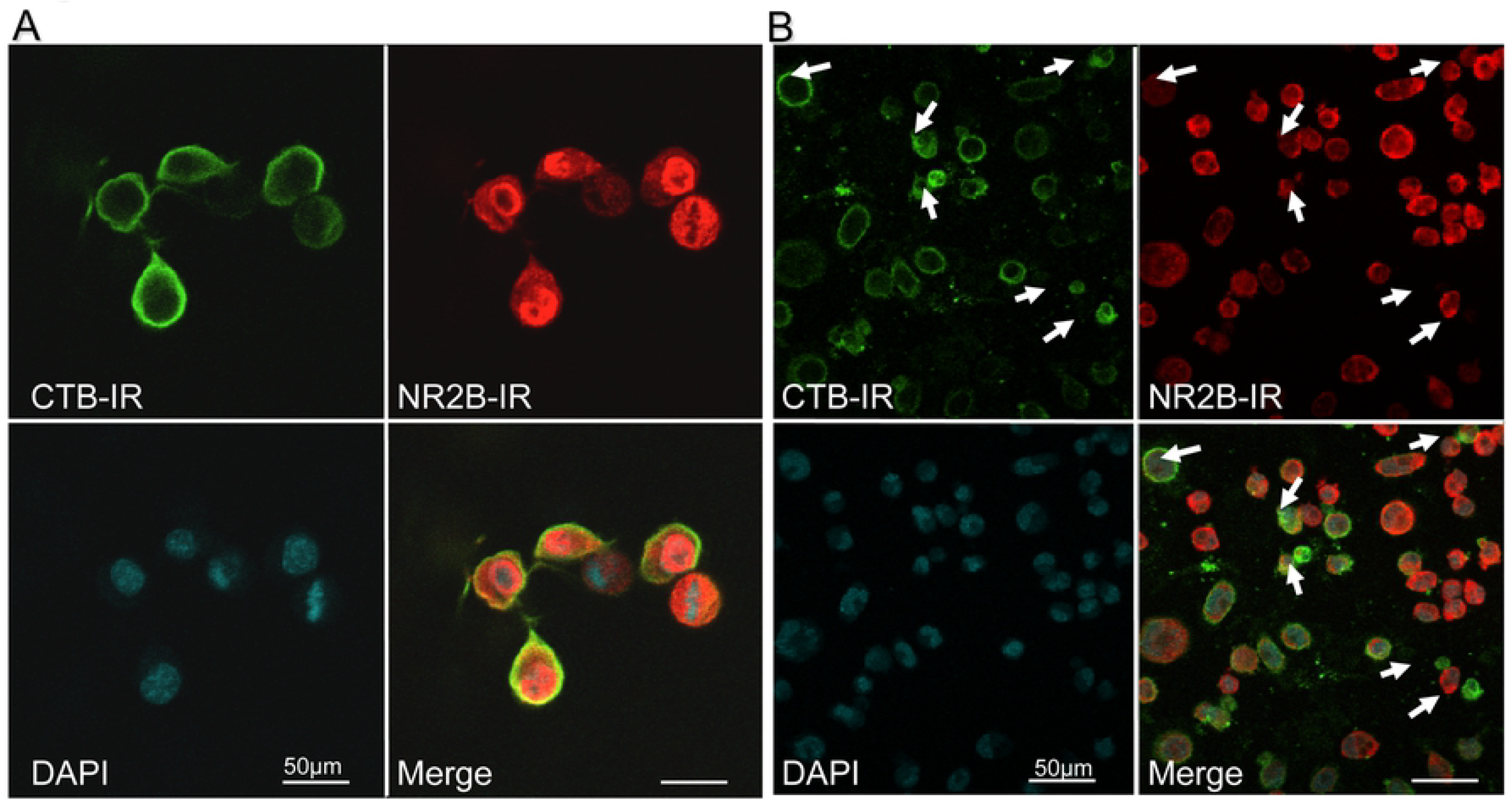
legend. Inhibition of NR2B immunoreactivity by CQ-dsNR2B complex. (A) Seven hours after incubation of Neuro2A cells with CQ-dsNR2B, uptake was evident as seen by positive staining of cells with CTB, but robust NR2B-IR was evident. (B) Two days after incubation of Neuro2A cells in medium containing CQ-dsNR2B, subsets of Neuro2A cells showed expression of CTB and markedly reduced expression of NR2B-IR, whereas cells that did not contain CTB showed robust NR2B-IR. Scale bar: 50μm

## Discussion

Here we show that the non-toxin subunit B of Cholera toxin, CTB that is routinely used as a neuroanatomical tracer can be used to deliver potentially therapeutic moieties to sub-populations of neurons. Given the selective uptake of CTB-dsRNA complex in cultured cells, this method has the potential to be a powerful way to study gene expression and cell signaling in a subpopulation of neurons, even in a mixed population of cells, and in their natural milieu. Currently, no such technique is available with this capacity.

Other toxins, peptides and receptors expressed on specific cell types or neurons have been used in past to deliver plasmid DNA to specific populations of cells or neurons; delivery of dsRNA (siRNA or miRNA) to neurons remains a challenge. Boulis’ group developed a peptide (Tet1) that is similar to tetanus toxin, is specifically taken up by motor neurons, and is retrogradely delivered to cell soma. Tet1-poly(ethylenimine) (Tet1-PEI) and neurotensin (NT)-PEI complexed with plasmid DNA have been evaluated as a neurontargeted delivery vehicle[20]. Plasmid DNA has also been conjugated to µ opioid receptor-liposome complexes for cell-specific delivery, but as is becoming apparent, chemical modification or fusion bacterial-mammalian proteins are immunogenic, and none of these methods can deliver unmodified nucleic acids or proteins.

Although here we focused on CTB as the carrier molecule, our method potentially allows us to encapsulate our resultant dsRNA/DNA nanoparticles with other carrier molecules such as isolectin B4 (IB4). IB4 is also used as a neuroanatomical tracer and targets a different population of neurons other than CTB. Potentially, IB4 and CTB can also be used in combination as carrier molecules to increase targeted populations of neurons. Consequently, our method has wide therapeutic potential.

The therapeutic areas that can be targeted with this delivery method could include any nervous system related disease. Additionally, since cholera toxin is a gut pathogen and enters the gut via epithelial cells expressing the cognate GM1 receptor, RNAi encapsulated in CTB can be targeted for gastrointestinal disorders.

This delivery platform can also serve as an alternative DNA and RNAi mammalian cell transfection reagents and vectors. Because only specific cell-types will take up the CTB-dsRNA or CTB-DNA complex, primary cultures of mixed populations can be transfected with much higher efficacy and efficiency. Moreover, since it is known that pure populations of neurons do not behave in the same manner as they do in their natural (*in vivo*) environment, this transfection method will allow us to target neurons in cultures that have both the neurons and the glia. Thus, functional studies in those targeted subpopulations of neurons can be performed.

## Acknowledgements

This work was supported by a CTSI pilot grant to AB. The authors wish to thank excellent technical help of Drs. Shilpi Mahajan, Pallavi Mhaske, and Ling-Hsuan Kung.

